# Central spindle microtubules are strongly coupled to chromosomes during both anaphase A and anaphase B

**DOI:** 10.1101/537290

**Authors:** Che-Hang Yu, Stefanie Redemann, Hai-Yin Wu, Robert Kiewisz, Tae Yeon Yoo, Reza Farhadifar, Thomas Müller-Reichert, Daniel Needleman

## Abstract

Spindle microtubules, whose dynamics vary over time and at different locations, cooperatively drive chromosome segregation. Measurements of microtubule dynamics and spindle ultrastructure can provide insight into the behaviors of microtubules, helping elucidate the mechanism of chromosome segregation. Much work has focused on the dynamics and organization of kinetochore microtubules, i.e. on the region between chromosomes and poles. In comparison, microtubules in the central spindle region, between segregating chromosomes, have been less thoroughly characterized. Here, we report measurements of the movement of central spindle microtubules during chromosome segregation in human mitotic spindles, and *Caenorhabditis elegans* mitotic and female meiotic spindles. We found that these central spindle microtubules slide apart at the same speed as chromosomes, even as chromosomes move towards spindle poles. In these systems, damaging central spindle microtubules by laser ablation caused an immediate and complete cessation of chromosome motion, suggesting a strong coupling between central spindle microtubules and chromosomes. Electron tomographic reconstruction revealed that the analyzed anaphase spindles all contain microtubules with both ends between segregating chromosomes. Our results provide new dynamical, functional, and ultrastructural characterizations of central spindle microtubules during chromosome segregation in diverse spindles, and suggest that central spindle microtubules and chromosomes are strongly coupled in anaphase.

## Introduction

Chromosome segregation is the essential biological processes, in which chromosomes are partitioned into the two daughter cells during cell division. In eukaryotes, chromosome segregation is carried out by the spindle, which is predominately composed of microtubules. Chromosomes first align in the middle of the spindle, and are then segregated in anaphase. It is often the case that chromosome segregation is accompanied by an elongation of the spindle. In many systems, the speed of spindle elongation is slower than chromosome segregation, such that chromosomes move closer to spindle poles over the course of anaphase. The movement of the chromosomes towards spindle poles is referred to as anaphase A; the elongation of the spindle (i.e. the separation of spindle poles) is referred to as anaphase B. The motion of anaphase A and anaphase B has been thought to depend on different behaviors of microtubules at different parts of the spindle (Asbury, 2017; Scholey et al., 2016).

The movement and turnover of spindle microtubules in anaphase have been measured with a variety of techniques, including photobleaching (Khodjakov et al., 2004; Mallavarapu et al., 1999; Redemann et al., 2017), photoconversion/photoactivation (Ferenz et al., 2010; Mitchison and Salmon, 1992; Vukusic et al., 2017) and fluorescence speckle microscopy (Brust-Mascher et al., 2004; Cameron et al., 2006; Pereira et al., 2016). Many previous studies have focused on characterizing kinetochore microtubule dynamics, providing great insights into the mechanisms of spindle assembly (Cameron et al., 2006; Ma et al., 2010; Maiato et al., 2005; Mitchison, 1989; Redemann et al., 2017), and the pole-ward chromosome motion in anaphase A and anaphase B (Brust-Mascher et al., 2004; LaFountain et al., 2004; Mitchison and Salmon, 1992). The behaviors of microtubules in the central spindle, between segregating chromosomes, have been less thoroughly characterized than kinetochore microtubules. It has been shown that microtubules of the central spindle slide apart while spindles elongate in diatoms, yeasts, flies, and human cells (Mallavarapu et al., 1999; Masuda et al., 1988; Saxton and McIntosh, 1987; Scholey et al., 2016; Vukusic et al., 2017). However, we are aware of few studies that *simultaneously* measure the speeds of central spindle microtubules and chromosomes (Vukusic et al., 2017) during Anaphase A and Anaphase B. Such data is potentially interesting, as it might provide further insight into the role of central spindle microtubules in chromosome segregation. Formally, there are three possibilities: central spindle microtubules might move slower, faster, or the same speed as segregating chromosomes (Figure 1).

**Figure 1.**
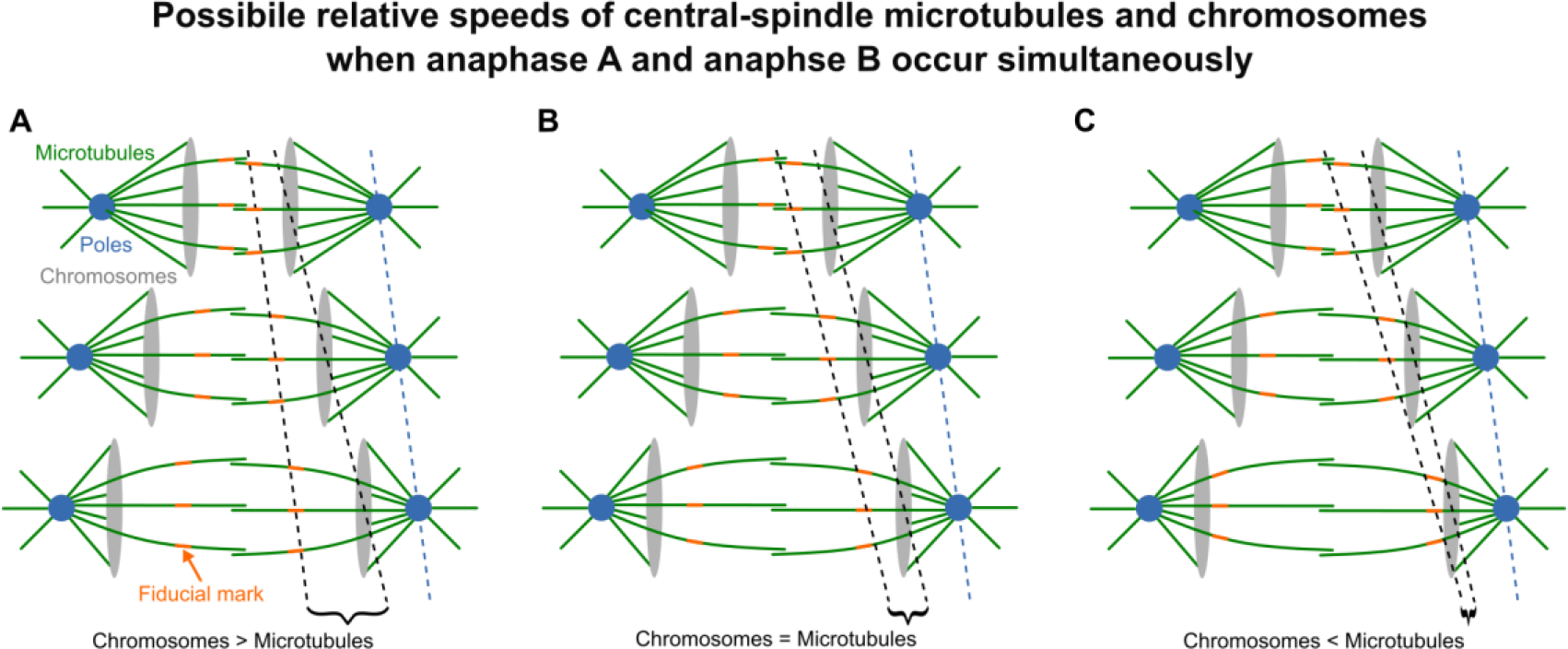
Possible relative speeds of central-spindle microtubules and chromosomes when anaphase A and anaphase B occur simultaneously. Central spindle microtubules might: (A) move slower than chromosomes; (B) move at the same speed as chromosomes; (C) move faster than chromosomes. For each case, the motion of a fiducial mark on central spindle microtubules (orange), chromosomes (gray), and spindle poles (blue), are indicated.

While simultaneous observation of microtubule and chromosome dynamics can reveal if their motions are correlated, perturbing spindle microtubules provides a means to test the extent to which their motions are coupled to the motion of chromosomes. A number of groups have used laser ablation to cut selected populations of microtubules and concluded that, in some anaphase spindles, central spindle microtubules resist pulling forces from astral microtubules (Aist and Bayles, 1991; Aist and Berns, 1981; Aist et al., 1993; Grill et al., 2001; Kronebusch and Borisy, 1981); in some anaphase spindles, central spindle microtubules exert pushing forces that contribute to anaphase chromosome motion (Khodjakov et al., 2004; Laband et al., 2017; Nahaboo et al., 2015; Vukusic et al., 2017). However, perturbation experiments must always be interpreted with care. One complication is the potential for redundant mechanisms. Thus, the observation that chromosome segregation can proceed without a connection to spindle poles in many systems (Elting et al., 2014; Khodjakov et al., 2004; Nahaboo et al., 2015; Nicklas, 1989; Vukusic et al., 2017) is equally consistent with two different explanations: that a loss of connection to the pole can be compensated by alternative processes, or that pole connections minimally contribute to chromosome motion. Another concern with any perturbation experiment is the possibility of collateral damage/off target effects. These concerns can be minimized (but not eliminated) by focusing on the immediate response to a rapid perturbation using a technique with a highly localized impact. In this regard, laser ablation with femtosecond pulses is advantageous, as cuts can be performed in seconds with ~300 nm resolution with no discernible impact outside of the ablated region (Schaffer et al., 2001a; Schaffer et al., 2001b; Vitek et al., 2010), unlike ablation with other lasers that are more typically used, which produce significant amounts of denatured protein (Brenner et al., 1980; Khodjakov et al., 2000; Khodjakov et al., 1997).

Light microscopy has insufficient resolution to resolve individual microtubules in spindles. Thus, a detailed study of spindle architecture requires alternative techniques, such as electron microscopy (Muller-Reichert et al., 2018). The structure of central spindle microtubules has been studied using electron microscopy in a number of spindles, which has revealed different structures in different spindles. In yeast and diatom spindles, central spindle microtubules span from the middle of the spindle to the poles (Ding et al., 1993; McDonald et al., 1977; Mcdonald et al., 1979; Winey et al., 1995; Winey et al., 2005). In contrast, in PtK1 mitotic spindles, many central spindle microtubules have both ends between chromosomes, while other central spindle microtubules extend from the inter-chromosomal region and terminate around kinetochore fibers (Mastronarde et al., 1993). In *C. elegans* female meiotic spindles, partial electron microscopy reconstruction revealed that many central spindle microtubules have both ends between chromosomes, some of which appear to make end-on contacts to chromosomes (Laband et al., 2017; Redemann et al., 2018). These different structures make it unclear to what extent central spindle microtubules are coupled to chromosomes in different spindles.

In this work, we investigated the relationship between central spindle microtubules and chromosomes during anaphase in human mitotic spindles, and *C. elegans* mitotic and female meiotic spindles. We used the same laser ablation and electron microscopy techniques on all of these spindles to avoid the potential complications of interpretation that can arise when different techniques are applied to different systems. We found that central microtubules slide apart at the same speed as chromosomes, even when chromosomes move closer to poles. Damaging the central spindle microtubules by laser ablation caused immediate and complete cessation of chromosome motion, even when Anaphase A and Anaphase B occur simultaneously, suggesting a strong coupling between central spindle microtubules and chromosomes. Our electron tomographic reconstructions further reveal that these anaphase spindles all contain microtubules with both ends between segregating sister chromosomes, and other central spindle microtubules which terminate on either chromosomes or kinetochore fibers. Taken together, this work suggests that central spindle microtubules are strongly coupled to chromosomes in anaphase spindles.

## Results

### Central-spindle microtubules slide apart at the speed of chromosome motion in anaphase human mitotic spindles

We set out to simultaneously measure the relative motion of central-spindle microtubules and chromosomes by generating a stable human cell line expressing mEOS3.2::tubulin and GFP::CENP-A. This cell line allowed us to photoconvert a subset of labeled microtubules between chromosomes while simultaneously tracking the motion of kinetochores and poles. We focused on studying chromosome motion 30-60 seconds after the onset of anaphase. Tracking the motions of kinetochores and spindle poles reveals that, at this time, chromosomes move away from the center of the spindle with a speed of 2.7 ± 0.2 μm/min, while poles move away from the spindle center at a speed of 1.5 ± 0.1 μm/min, so chromosomes move toward poles at a speed of 1.2 ± 0.2 μm/min (n=8; Figure 2A). Thus, our experiments probe a time window in which the speed of chromosome-to-pole motion is similar to the speed of pole motion. Next, a line of fluorescence labeled microtubules across the spindle was photoconverted 30-60 seconds after the onset of anaphase. The single line of photoconverted tubulin split apart over time, demonstrating the sliding of central-spindle microtubules (Figures 2B, magenta; Movie S1). Kymographs revealed that the speed of central-spindle microtubule sliding was similar to the speed of chromosome motion, and substantially faster than pole motion (Figure 2C). To quantify the speed of the central-spindle microtubules, we took line profiles and tracked the motion of photoactivated regions (Figure 2D), in addition to chromosomes and poles (see methods). We found that the photoconverted microtubules slide at a speed of 2.8 ± 0.1 μm/min (n=46), which is indistinguishable from the speed of chromosome motion, 2.7 ± 0.1 μm/min, (n=46, p = 0.85) and substantially greater than the speed of pole motion, 1.5 ± 0.1 μm/min (n=46, p<10^−11^) (Figure 2D, right). This result shows that central-spindle microtubules and chromosomes move at the same speed in human mitotic anaphase spindles, even at times when anaphase A and anaphase B occur simultaneously, suggesting a coupling between them.

**Figure 2.**
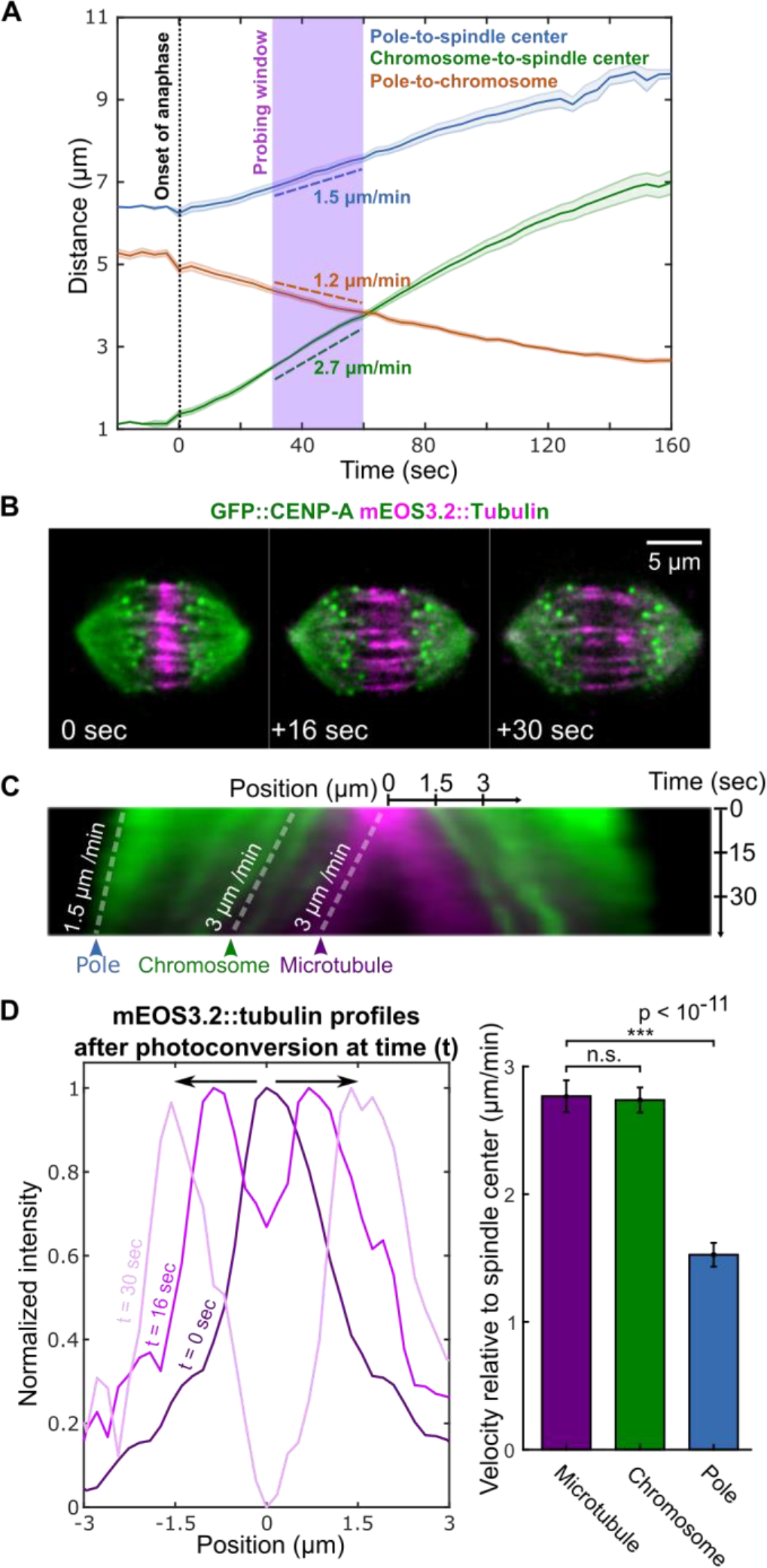
Central-spindle microtubules move apart at the same speed as chromosomes in human mitotic spindles. (A) Change of pole-to-spindle center (blue), chromosome-to-spindle center (green), and chromosome-to-pole (orange) distances during chromosome segregation. The slopes show the averaged velocity of above three quantities during the probing window (purple), in which the experiments and observations were conducted. (B) Time-lapse images of GFP::CENP-A (green) and mEOS3.2::tubulin (green before photoconversion; magenta after photoconversion) human mitotic spindles with photoconverted microtubules between chromosomes. Time zero is the onset of photoconversion. (C) A kymograph of the spindle shown in (B), with white dashed lines illustrating that the photoconverted microtubules and chromosomes move apart at the same speed, and both are faster than poles. (D) Line-profiles of photoconverted mEOS3.2-tubulin between chromosomes after photoconversion (left) from the kymograph in (C), with arrows to indicate the split of the photoconverted region. Bar plot of velocities of microtubules (magenta), chromosomes (green) and spindle poles (blue) in human mitotic spindles 30-60 seconds after the onset of anaphase (right). Error bars are SEMs (n.s., not significant).

### Laser-ablating central-spindle microtubules immediately stops anaphase chromosome motion in human mitotic spindles

We next sought to investigate to what extent central-spindle microtubules are coupled to chromosome motion in human mitotic spindles. We built a custom laser ablation system utilizing an ultrafast femtosecond laser, which enables sub-diffraction-limited cuts (Vogel et al., 2005) in nearly arbitrary three-dimensional patterns. These cuts are performed within a few seconds and generate minimal collateral damage outside of the ablated region (see methods). We performed a rectangular plane-cut, 12-μm in length by 6-μm in depth, perpendicular to the spindle axis between separating chromosomes 30-60 seconds after the onset of anaphase (Figure 3A; left; Movie S2). This ablation led to an immediate cessation of chromosome motion, reducing its speed to 0.1 ± 0.3 μm/min (n=6; indistinguishable from zero, p = 0.83). After approximately ~20 seconds, chromosome segregation resumed with a speed similar to controls, presumably due to the microtubule re-growth, replacing the damaged population of microtubules (Figure 3A, right). These results further argue that central-spindle microtubules are strongly coupled to chromosome motion, even at times when anaphase A and anaphase B occur simultaneously.

**Figure 3.**
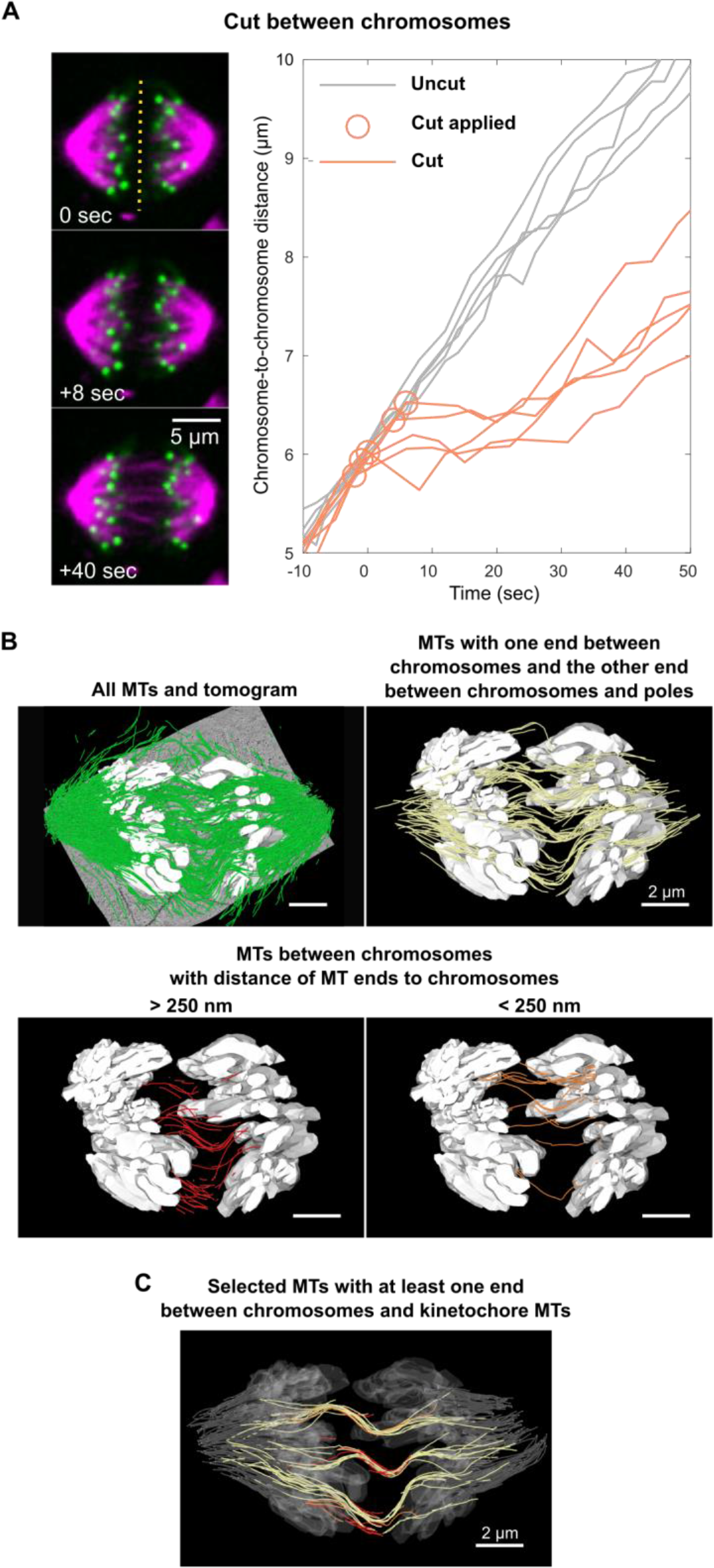
The functional and structural characterization of central-spindle microtubules in human mitotic spindles. (A) Time-lapse images of GFP::CENP-A (green) and mCherry::tubulin (magenta) human mitotic spindles with laser ablation of microtubules between chromosomes (left). Dotted lines indicate the timing and location of laser ablation. Curves of inter-chromosome distance versus time with laser ablation of microtubules between chromosomes (orange) with uncut spindles (gray) for reference (right). (B) Electron tomographic reconstruction of microtubules in human mitotic spindles, showing all microtubules (green, upper left) overlaid on the tomogram. Some microtubules have both ends between chromosomes, with neither (red, lower left) or either one (orange, lower right) of their ends contacting chromosomes; other microtubules have only one end between chromosomes (yellow, upper right). (C) Electron tomographic reconstruction of selective microtubule bundles consisting of the three classes of microtubules (red, orange, and yellow) in (B), kinetochore microtubules (gray), and chromosomes (translucent).

### Electron tomography reveals the presence of inter-chromosomal microtubules in human mitotic spindles

To further investigate how the central-spindle microtubules are coupled with chromosomes, we next studied the structure of human mitotic spindles in anaphase using large-scale electron tomographic reconstructions (Figure 3B, green). This work revealed the presence of inter-chromosomal microtubules: some of which had two free ends between chromosomes (Figure 3B, red), others of which had one end near the center of the spindle with the other end contacting chromosomes (Figure 3B, orange). Other microtubules extended from the inter-chromosomal region, passed chromosomes, reaching a micron or so beyond them (Figure 3B, yellow). We found no microtubules, which extended all the way from the pole to the region between chromosomes, and no microtubules, which passed across the entire inter-chromosomal region, directly bridging microtubules between the two poles. The three classes of microtubules we observed are often tightly associated into bundles, which appear to connect inter-chromosomal microtubules to kinetochore microtubules, near chromosomes (Figure 3C). This structure illustrates how inter-chromosomal microtubules in the central-spindle region may cross-link to kinetochore microtubules, which thus would couple the motion of central-spindle microtubules to the motion of chromosomes. This structure of central spindle microtubules is reminiscent of what was previously seen in PtK1 cells (Mastronarde et al., 1993).

### Central-spindle microtubules slide apart at the speed of chromosome motion in *C. elegans* anaphase mitotic spindles

Having characterized dynamics, function, and structure of central-spindle microtubules in human mitotic spindles, we next explored if central-spindle microtubules in other spindles have similar behavior and organization. We first set out to measure the relative motion of central-spindle microtubules and chromosomes in anaphase *C. elegans* mitotic spindles expressing GFP::tubulin and mCherry::histone. We were unable to reliably photobleach central spindle microtubules in *C. elegans* mitotic spindles due to the rapid oscillations of these spindles in anaphase (Movie S3). To better maintain the spindle in the focal plane, we knocked down GPR-1/2 using RNA interference (RNAi), which greatly reduce the cortically pulling forces (Movie S4) (Grill et al., 2003; Park and Rose, 2008; Srinivasan et al., 2003). To understand the effect of *gpr-1/2(RNAi*) on chromosome segregation, we acquired time-lapse images of spindles for the first 3 minutes of anaphase in the first mitotic division of wildtype and *gpr-1/2(RNAi) C. elegans* embryos (Figure 4A; Movie S3 and S4), and averaged the traces of pole motion, chromosome motion, and chromosome-to-pole motion (Figure 4B). In the wildtype embryos, the chromosome-to-pole distance stayed almost constant during this time (Figure 4B, upper, orange) (Oegema et al., 2001): i.e. the mean speed of chromosomes, 1.1 ± 0.02 μm/min, was indistinguishable from the mean speed of poles, 1.1 ± 0.05 μm/min (Figure 4C, upper, n=5, p = 0.93). In contrast, time-lapse microscopy of *gpr-1/2(RNAi)* embryos showed that while pole motion was significantly reduced, the speed and extent of chromosome motion remained very similar to that in controls (compare Figure 4A upper, lower; Movie S3 and S4). We averaged the traces of pole motion, chromosome motion, and chromosome-to-pole motion, and found that the distance between chromosomes and poles continuously decreased over the course of anaphase (Figure 4B, lower, orange), as the mean speed of poles, 0.45 ± 0.04 μm/min, was significantly less than the mean speed of chromosomes, 0.90 ± 0.03 μm/min (Figure 4C, lower, n=8, p < 10^−6^). Consistent with previous results (Nahaboo et al., 2015), this experiment demonstrates that cortically pulling forces are not required for chromosome segregation in *C. elegans* mitosis. Since GPR-1/2 knockdown greatly reduced spindle movement and only had a minor impact on chromosome motion, we used this RNAi condition to investigate the relative motion of chromosomes and central-spindle microtubules.

**Figure 4.**
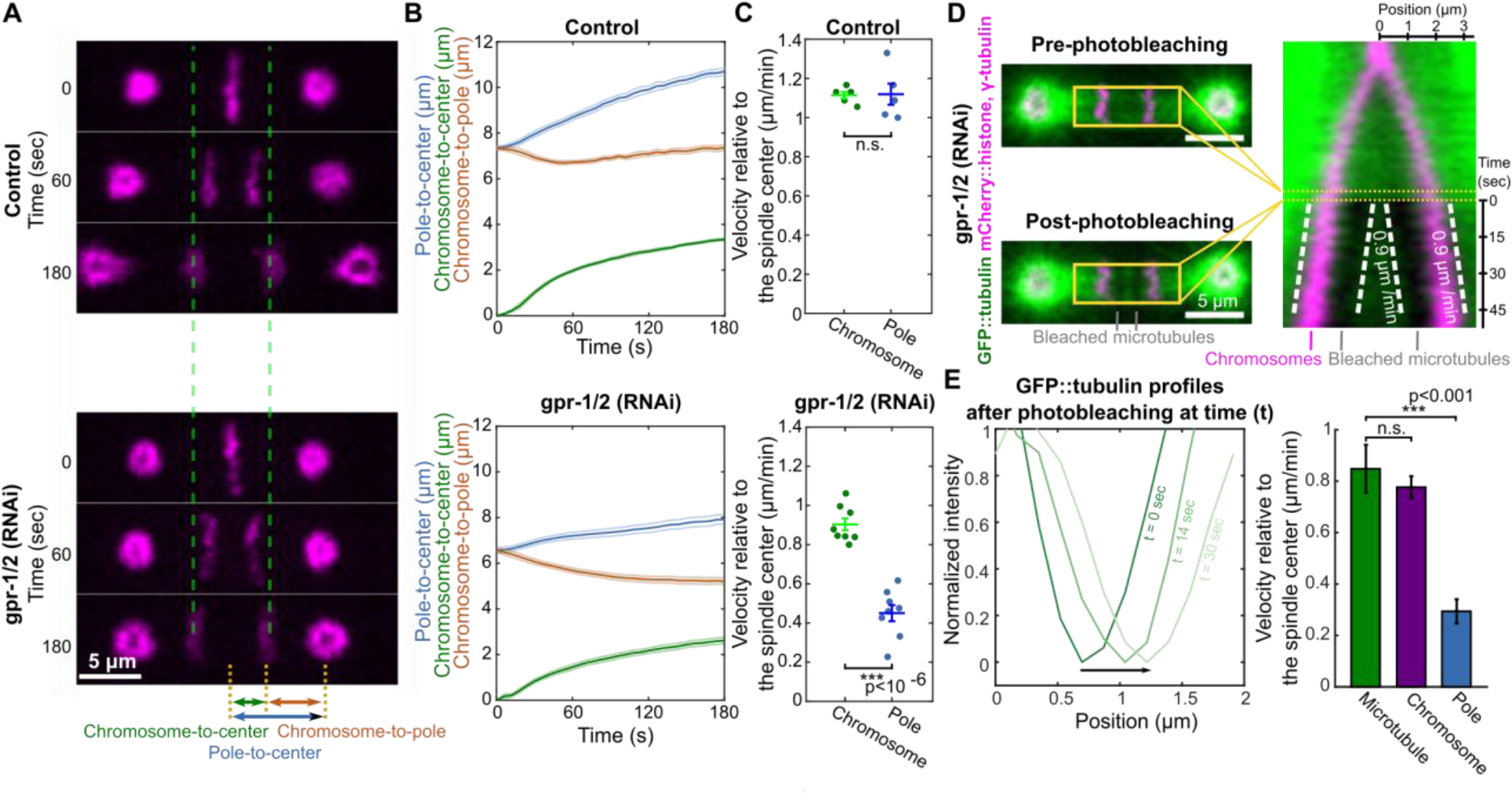
Central-spindle microtubules move apart at the same speed as chromosomes in *C. elegans* mitotic spindles. (A) Time-lapse images of mCherry::histone and mCherry::γ-tubulins (magenta) in control (upper) and *gpr-1/2 (RNAi)* spindles (lower). Time zero is the onset of anaphase. (B) Averaged pole-to-spindle center distance (blue), chromosome-to-spindle center distance (green), and chromosome-to-pole distance (orange) in control (upper, n=5) and *gpr-1/2 (RNAi)* (lower, n=8) embryos. (C) Scatter plots of velocities of chromosomes (green) and spindle poles (blue) in control (upper, n=5) and *gpr-1/2 (RNAi)* (lower, n=8) embryos during the first 3 minutes of anaphase. Error bars are SEMs (n.s., not significant). (D) Images of a GFP::tubulin (green), mCherry::histone and mCherry::γ-tubulin (magenta) spindle before and after photobleaching, indicting the location of photobleached regions (left). A kymograph from the indicated region, with white dashed lines illustrating that the bleached regions and chromosomes move apart at the same speed (right); time zero is the onset of photobleaching. (E) Line-profiles of GFP-tubulins (left) over the region corresponding to the kymograph in (D), with an arrow to indicate the shift of the photobleached region. Bar plot of velocities of microtubules (green), chromosomes (purple) and spindle poles (blue) in *gpr-1/2 (RNAi)* embryos 40-60 seconds after the onset of anaphase (right). Error bars are SEMs (n.s., not significant).

We conducted photobleaching experiments in *gpr-1/2(RNAi)* embryos 40-60 seconds after the onset of anaphase, when the chromosome-to-pole speed was 0.5 ± 0.07 μm/min (n=7) and the pole-to-spindle center speed was 0.3 ± 0.05 μm/min (n=7). Thus, at this time, the chromosome-to-pole distance shrank at a faster rate than the poles separate (i.e. Anaphase A and Anaphase B were both underway at this time). We photobleached GFP-labeled tubulin to create two parallel fiducial marks on microtubules near the spindle center (Figure 4D, left). Kymographs showed that these fiducial marks moved apart at a speed similar to the speed of chromosomes separation (Figure 4D, right). The movement of the photobleached marks demonstrates that that microtubules between chromosomes slide apart in anaphase. To further quantify this motion, we took line-profiles of the bleached region (Figure 4E), tracked their motion and the motion of chromosomes (see methods), and found that the central-spindle microtubules move at a speed of 0.8 ± 0.09 μm/min (n=7), which is indistinguishable from the speed of chromosome movement (0.8 ± 0.04 μm/min, n=7, p = 0.53), and substantially greater than the speed of pole movement (0.3 ± 0.05 μm/min, n=7, p < 0.001). Therefore, our results reveal that central-spindle microtubules slide apart at the same speed as chromosomes, even at times when anaphase A and anaphase B occur simultaneously. Similar to the results in the human mitotic spindles, these results argue for a coupling between the central-spindle microtubules and chromosomes in *C. elegans* mitotic spindles.

### Laser-ablating central-spindle microtubules immediately stops anaphase chromosome motion in *C. elegans* mitotic spindles

We next sought to investigate the coupling between the central-spindle microtubules and chromosomes in *C. elegans* mitotic spindles using our custom laser ablation system. We first cut a rectangular plane, 4-μm in length by 6-μm in depth, perpendicular to the spindle axis halfway-through between separating chromosomes in *gpr-1/2(RNAi)* embryos (Figure 5A; Movie S5). Cuts were performed 30-60 seconds after the onset of anaphase when chromosomes and poles move away from the spindle center at a speed of 1.2 ± 0.1 μm/min and 0.5 ± 0.1 μm/min, respectively, so that chromosomes move toward poles at a speed of 0.7 ± 0.1 μm/min. Thus, at this time, the chromosome-to-pole distance shrinks at a faster rate than the poles separate (i.e. Anaphase A and Anaphase B were both underway at this time). The ablation led to an immediate cessation of chromosome motion, reducing its speed to 0.1 ± 0.2 μm/min (n=7; indistinguishable from zero, p = 0.73). After approximately ~20 seconds, chromosome segregation resumed with a speed similar to the uncut controls, presumably due to microtubule re-growth and replacement of the damaged microtubule population (Figure 5A, right). Thus, ablating microtubules between chromosomes completely stopped chromosome motion, while, at the time of these cuts, chromosome-to-pole distance was decreasing and spindle poles were separating. The result further argue that the central-spindle microtubules are strongly coupled to chromosome motion in *gpr-1/2(RNAi) C. elegans* embryos.

**Figure 5.**
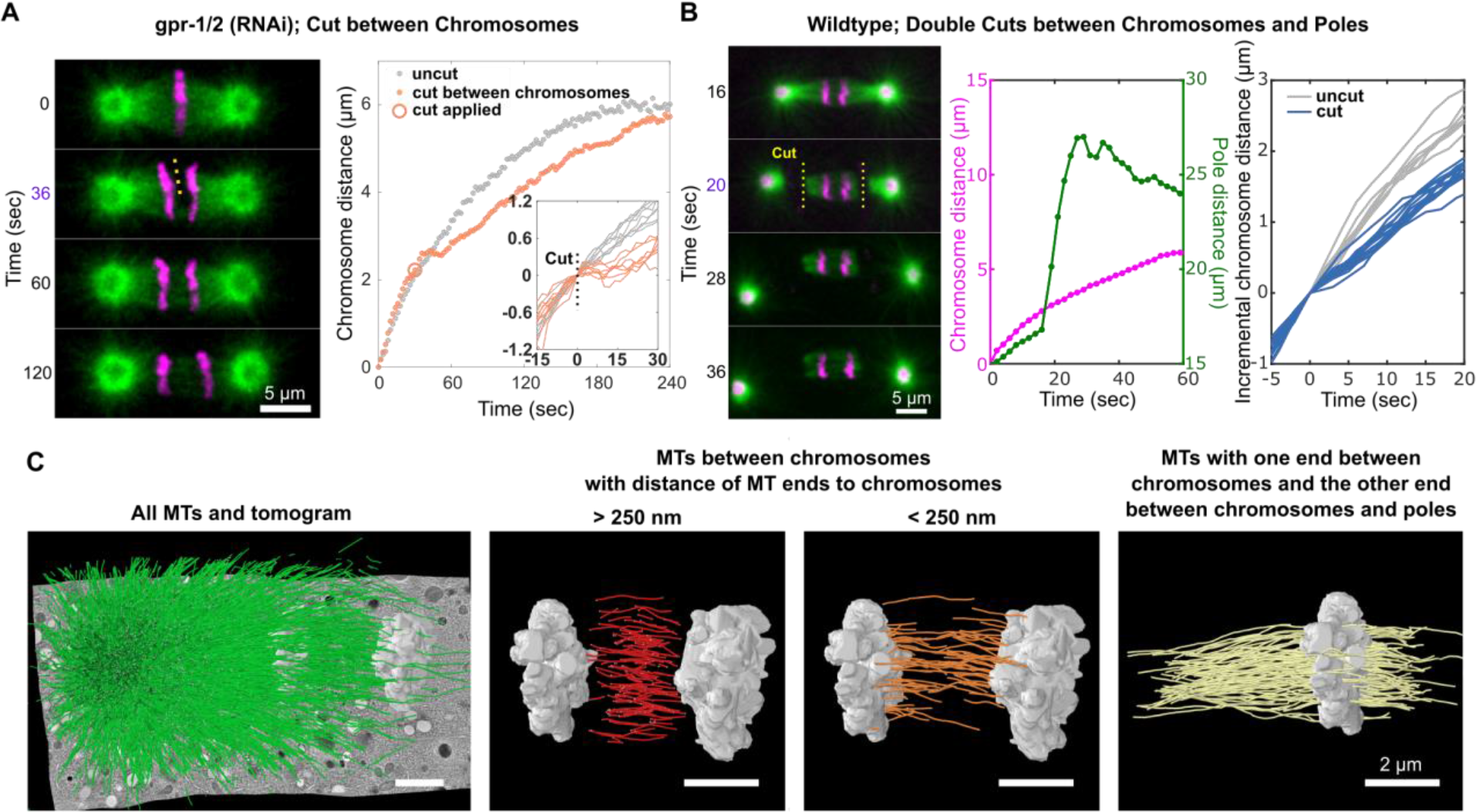
The functional and structural characterization of central-spindle microtubules in *C. elegans* mitotic spindles. (A) Time-lapse images of GFP::tubulin (green) and mCherry::histone (magenta) in *gpr-1/2 (RNAi) C. elegans* spindles (left) when microtubules are cut between chromosomes. Time zero is the onset of anaphase. Dotted lines indicate the timing and location of laser ablation. Corresponding plots (right) of chromosome distance as a function of time, with an uncut spindle (grey) for reference. Example plots (right, inserts) of the change in chromosome distance after the cut from multiple spindles, aligned relative to the timing of the cut, with uncut spindles (gray) for reference. **(B)** Time-lapse images of GFP::tubulin (green) and mCherry::histone, γ-tubulin (magenta) in wildtype *C. elegans* spindles when microtubules are cut between chromosomes and poles on both sides of the spindle (left). Time zero is the onset of anaphase. Yellow dotted lines indicate the timing and location of laser ablation. Corresponding traces of chromosome-to-chromosome distance and pole-to-pole distance to the time-lapse image as a function of time (middle). Example plots of the change in chromosome distance after the cut from multiple spindles, aligned relative to the timing of the cut, with uncut spindles (gray) for reference (right). (C) Electron tomographic reconstruction showing all traced microtubules (green) overlaid on a tomogram; some microtubules have both ends between chromosomes, with neither end (red) or either one (orange) contacting chromosomes; other microtubules have only one end between chromosomes (yellow, all such microtubules in a half-spindle).

This strong coupling in the embryos with GPR-1/2 depleted might be due to an activation of compensatory processes not present in wildtype embryos. To test the coupling between central-spindle microtubules and chromosomes in the wildtype embryos, we ablated two rectangular regions, 8-μm in length by 6-μm in depth, between chromosomes and spindle poles on both sides of the spindle (Figure 5B, left) while the sister chromosomes were moving apart. After the cuts, the spindle poles detached from the central body of the spindle (Figure 5B, middle, green) and rapidly moved apart at a speed of 60.8 ± 2.2 μm/min (n=13). In the subsequent ~15 seconds the poles underwent irregular movements while the central body of the spindle showed no correlated displacement or rotation, further arguing that the cut disconnected the central body of the spindle from the poles (Movie S6). Chromosome segregation continued after the cuts (Figure 5B, middle, magenta). The cuts did not pause chromosome motion, but reduced their speed from 9.2 ± 0.3 μm/min (n=7) in controls, to 5.2 ± 0.1 μm/min (n=13, p = < 10^−10^) (Figure 5B, right). Thus, while previous work showed that *C. elegans* mitotic spindles can segregate chromosomes if the poles are destroyed before the onset of anaphase (Nahaboo et al., 2015), our new results demonstrate that chromosomes can continue segregating even immediately after severing the connection between chromosomes and poles in anaphase. The continued motion of chromosomes after sudden detachment from both spindle poles argues that, even in wildtype *C. elegans* mitotic embryos, the central-spindle microtubules are coupled to chromosomes in a manner that is not entirely dependent on connections through spindle poles.

### The anaphase *C. elegans* mitotic spindle contains inter-chromosomal microtubules

To further investigate how the central-spindle microtubules couple to chromosomes, we performed serial-section electron tomography to reconstruct the *C. elegans* anaphase mitotic spindles (Figure 5C, left). Our reconstructions revealed the presence of inter-chromosomal microtubules: some of which had two free ends between chromosomes (Figure 5C, red), others of which had one end near the center of the spindle with the other end contacting chromosomes (Figure 5C, orange). In addition, there are also microtubules with one end extending from the inter-chromosomal region passing around chromosomes to contact kinetochore microtubules (Figure 5C, yellow). Again, we also found no microtubules, which extended all the way from the pole to the region between chromosomes, and no microtubules, which passed across the entire inter-chromosomal region, directly bridging microtubules between the two poles. Therefore, electron tomography demonstrates that inter-chromosomal microtubules also exist in anaphase *C. elegans* mitotic spindles, similar to those in human anaphase spindles. The structure suggests that the coupling from central-spindle microtubules to chromosomes could either through direct contact at the inner surface of chromosomes (Figure 5C, orange microtubules) from the side of the central-spindle region, or, alternatively, the connection to chromosomes could be indirectly, by microtubules with one end extending from the inter-chromosomal region passing around chromosomes to contact kinetochore microtubules (Figure 5C, yellow). The lack of microtubules connecting the central-spindle region directly to the poles may explain the continued motion of chromosomes immediately after severing the connections between poles and chromosomes in anaphase.

### Electron tomography confirms the presence of inter-chromosomal microtubules in *C. elegans* female meiosis

Having characterized the dynamics, function, and structure of central-spindle microtubules in human and *C. elegans* mitotic spindles, we next sought to apply the same approach to study a spindle with very different morphology and dynamics: acentrosomal *C. elegans* female meiotic spindles. We first generated electron tomographic reconstructions of all microtubules in *C. elegans* female meiotic spindles at various stages of anaphase (Figure 6A, left). Nearly all microtubules in the spindle lie between chromosomes at these stages of anaphase (Figure 6A, left). The organization of inter-chromosomal microtubules in *C. elegans* female meiotic spindles is highly reminiscent of those found in *C. elegans* mitotic spindles: some microtubules have both their ends between the chromosomes, without touching the chromosomes (Figure 6A, center), while some microtubules make end-on contact with chromosomes, with their other end terminating between the chromosomes (Figure 6A, right). Our observation of numerous inter-chromosomal microtubules in anaphase of *C. elegans* female meiotic spindles is consistent with previous partial electron tomography reconstructions (Laband et al., 2017).

**Figure 6.**
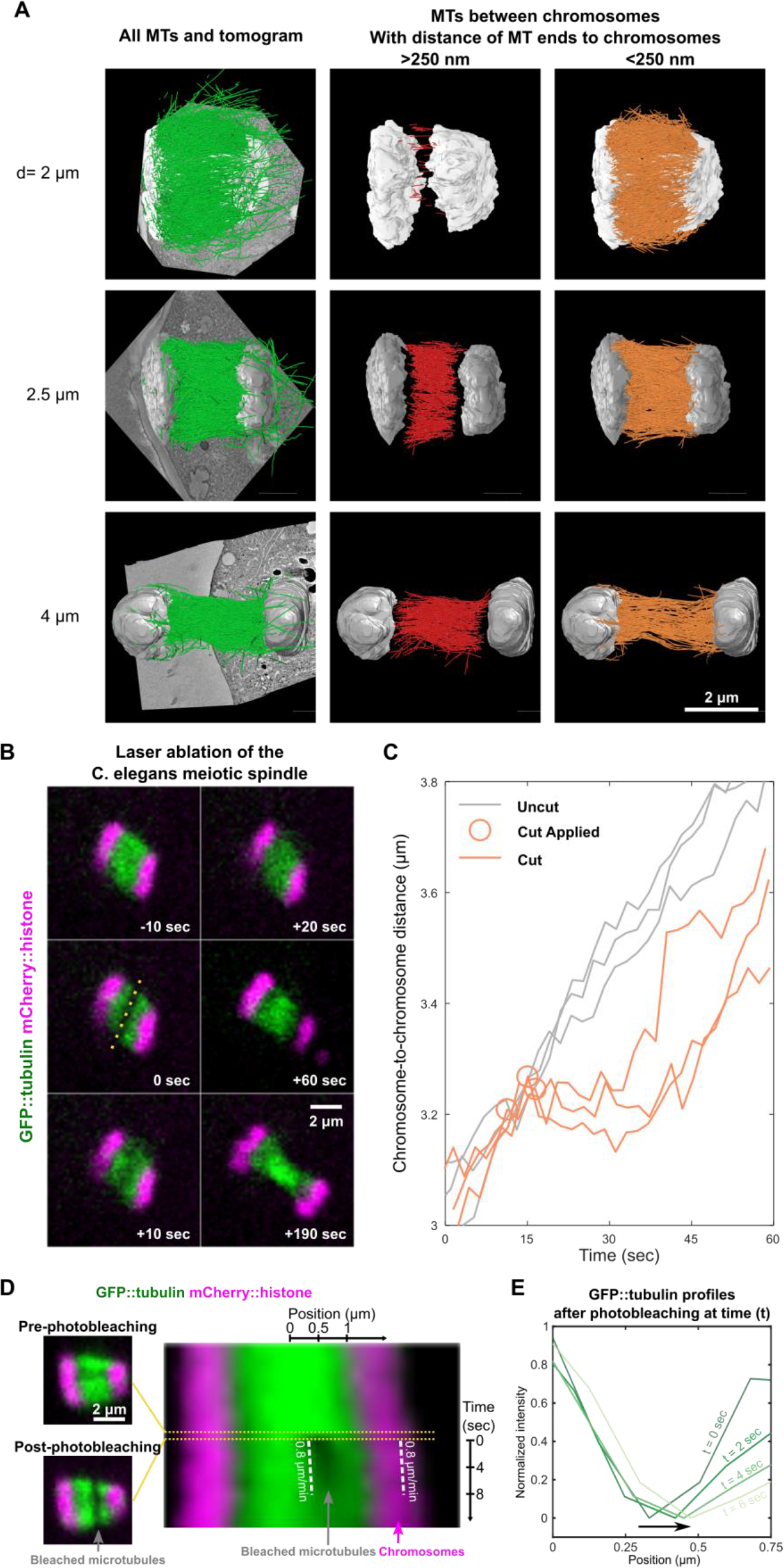
The functional, structural, and dynamical characterization of inter-chromosomal microtubules in *C. elegans* female meiotic spindles. (A) Electron tomographic reconstructions of microtubules in *C. elegans* meiotic spindles with inter-chromosomal distances (d) of 2 μm (upper row), 2.5 μm (middle row), and 4 μm (lower row), showing all microtubules (green) overlaid on tomograms. Some microtubules have both ends between chromosomes, with neither (red) or either one (orange) of their ends contacting chromosomes. (B) Time-lapse images of a GFP::tubulin (green) and mCherry::histone (magenta) *C. elegans* meiotic spindle, with the dotted line indicating the timing and location of laser ablation. (C) Inter-chromosomal distance versus time for spindles cut between chromosomes (orange), with uncut spindles (grey) for reference. (D) Images of a GFP::tubulin (green) and mCherry::γ-tubulin (magenta) spindle before and after photonbleaching (left), showing the position of the photobleached region (left). A kymograph of a spindle, with white dashed lines illustrating that the bleached regions and chromosomes move apart at the same speed; time zero is the onset of photobleaching (right). (E) Line-profiles of GFP-tubulin over the region corresponding to the kymograph in (D), with an arrow to indicate the shift of the bleached center.

### Laser-ablating central-spindle microtubules immediately stops anaphase chromosome motion in *C. elegans* meiotic spindles

To test the coupling of these inter-chromosomal microtubules to chromosomes, we used laser ablation to cut them during anaphase (Figure 6B; Movie S7). We found that, as in mitotic spindles, this damage caused an immediate cessation of chromosome motion, with their speed reducing to 0.01 ± 0.01 μm/min (n=6; indistinguishable from zero, p=0.36), from a speed of 0.7 ± 0.1 μm/min (n=5) in controls (Figure 6C). Similar to mitotic spindles, chromosome motion resumed at a speed similar to controls after approximately 20 seconds, presumably due to the spontaneous repair of the damaged population of microtubules. Thus, this result demonstrates that inter-chromosomal microtubules are strongly coupled to chromosomes in *C. elegans* female meiotic spindles at the stages we studied. A previous laser ablation study also found that cutting inter-chromosomal microtubules halted chromosome motion in *C. elegans* meiotic spindles (Laband et al., 2017), though, in that study severed spindles never recovered, likely due to severe collateral damage caused by using a UV laser for ablation.

### Inter-chromosomal microtubules slide apart at the speed of chromosome motion in *C. elegans* female meiotic spindles

We next investigated the relative motion of inter-chromosomal microtubules and chromosomes in *C. elegans* female meiotic spindles (Figure 1). We photobleached GFP-labeled tubulin to create a fiducial mark on inter-chromosomal microtubules adjacent to a set of sister chromosomes during anaphase and found that they moved at a similar speed as chromosomes moved (Figure 6D). To better quantify this motion, we made line-profiles of the photobleached regions and tracked their motion, along with the motion of chromosomes (Figure 6E) (see methods). Averaging measurements over multiple spindles gave a mean speed of microtubule sliding away from the spindle center of 0.5 ± 0.2 μm/min (n=6), indistinguishable from the mean speed of chromosome movement, 0.3 ± 0.1 μm/min (n=6, p=0.3). These results further argue that central spindle microtubules are strongly coupled to anaphase chromosomes in *C. elegans* female meiotic spindles.

## Discussion

We used optical microscopy, laser ablation, and electron tomography to characterize central spindle microtubules in anaphase in human mitotic spindles, and *C. elegans* mitotic and female meiotic spindles. We found that central spindle microtubules slide apart at the same speed as chromosomes, even at times when Anaphase A and Anaphase B occur simultaneously. Laser ablating central spindle microtubules lead to an immediate cessation of chromosome motion, even at times when Anaphase A and Anaphase B occur simultaneously. These results argue that central spindle microtubules are tightly coupled to chromosomes in anaphase. Because of technical limitations, our most detailed characterization of *C. elegans* mitotic spindles were performed when pulling forces were inhibited by *gpr-1/2(RNAi)*, a perturbation that only minimally impacted chromosome motion (while greatly reducing pole oscillations and movements). Our laser ablation experiments demonstrate that chromosomes continue to separate even immediately after severing the connection between chromosomes and poles, suggesting that a tight coupling between central spindle microtubules and chromosomes is also present in wildtype *C. elegans* mitotic spindles.

To gain further insight into the manner by which central spindle microtubules are couples to chromosomes in anaphase, we also performed large-scale electron tomography reconstructions of these microtubules in human mitotic spindles, and *C. elegans* mitotic and female meiotic spindles. We found that all of these spindles contained microtubules with both ends between segregating chromosomes, some of which appeared to directly contact chromosomes. Human and *C. elegans* mitotic spindles both also contained microtubules with one end extending from the inter-chromosomal to contact kinetochore microtubules. In all the spindles we studied, we found no microtubules, which extended all the way from the pole to the region between chromosomes, and no microtubules, which passed across the entire inter-chromosomal region, directly bridging kinetochore-microtubules from the opposite two poles. Thus, our data argues that central spindle microtubules could be connected directly to chromosomes, or the coupling could be indirect through their mutual connection to kinetochore microtubules (or both). It is an exciting challenge for future research to disentangle these possibilities.

## Methods

### *C. elegans* strains

Strain SA250 (tjIs54 [pie-1p::GFP::tbb-2 + pie-1p::2xmCherry::tbg-1 + unc-119(+)]; tjIs57 [pie-1p::mCherry::his-48 + unc-119(+)]), and a strain expressing GFP::tubulin and mCherry::histone (a gift from Marie Delattre lab) were used for experiments of fluorescence imaging, laser ablation, and fluoresce recovery after photobleaching. Wildtype (N2) *C. elegans* embryos and oocytes were used for the preparation of electron tomography. All strains were cultured at 24°C and fed on OP50 bacteria on nematode growth medium plates.

### Human cell lines and culture

Human Bone Osteosarcoma Epithelial (U2OS) cells were engineered to stably express fluorescence-labeled proteins by retroviral transfections. Three stable fluorescence U2OS lines were used: one expresses GFP::CENP-A, and mCherry::tubulin; another expresses GFP::centrin, GFP::CENP-A, and mCherry::tubulin; the other expresses GFP::CENP-A, and mEOS3.2::tubulin U2OS cell lines were maintained in Dulbecco’s modified Eagle’s medium (DMEM, Gibco) supplemented with 10% Fetal Bovine Serum (FBS, Gibco), and 50 IU/ml penicillin and 50 μg/ml streptomycin (Gibco) at 37°C in a humidified atmosphere with 5% CO2.

### *C. elegans* imaging preparation and RNA interference

For imaging of mitotic spindles, gravid *C. elegans* hermaphrodites were cut in half, and the released embryos were transferred onto a 4% agarose pad between a slide and a coverslip (Walston and Hardin, 2010). Meiotic spindles in oocytes were observed in uterus, as adult hermaphrodites were mounted between a coverslip and a thin 4% agarose pad on a slide. Polystyrene microspheres (Microspheres 0.10 μm, Polysciences, Inc.) in solution were added to help immobilize worms.

RNA interference (RNAi) was carried out following the RNAi feeding protocol from the Ahringer lab (Kamath et al., 2001). Plasmid pMD082 (a gift from Marie Delattre lab) containing the sequence of *gpr-1/2* was cloned into the L4440 plasmid, which was transformed into HT115 bacteria. L2 hermaphrodites were transferred to *gpr-1/2 (RNAi)* plates and fed on the RNAi bacterial lawn for 36–48 h at 24°C.

### Human cell imaging preparation

In preparation for imaging, cells were grown on a 25-mm diameter, #1.5-thickness, round coverglass coated with poly-D-lysine (GG-25-1.5-pdl, neuVitro) to 80~90% confluency. The cells were incubated in imaging media, which is FluoroBrite™ DMEM (Gibco) supplemented with 4mM L-glutamine (Gibco) and 10mM HEPES, for 15~30 minutes before imaging. The coverglass was mounted on a custom-built temperature controlled microscope chamber at 37°C, while covered with 1.5 ml of imaging media and 2 ml of white mineral oil (VWR). An objective heater (Bioptech) was used to maintain the objective at 37°C.

### Sample preparation for electron tomography

Wild-type (N2) *C. elegans* embryos and oocytes collected in cellulose capillary tubes (Pelletier et al., 2006; Redemann et al., 2018) and HeLa Kyoto cells grown on sapphire discs (Guizetti et al., 2011) were high-pressure frozen as described using an either an EM PACT2+RTS or an EM ICE high-pressure freezer (Leica Microsystems, Vienna, Austria)(Muller-Reichert et al., 2007). Freeze substitution was performed over 2-3 d at −90 °C in anhydrous acetone containing 1 % OsO4 and 0.1 % uranyl acetate. Samples were embedded in Epon/Araldite and polymerized for 2-3 d at 60 °C. Serial semi-thick sections (300 nm) were cut using an Ultracut UCT Microtome (Leica Microsystems, Vienna, Austria), collected on Formvar-coated copper slot grids and post-stained with 2 % uranyl acetate in 70 % methanol followed by Reynold’s lead citrate. For dual-axis electron tomography (Mastronarde, 1997) series of tilted views were recorded using a TECNAI F30 transmission electron microscope (FEI Company, Eindhoven, The Netherlands) operated at 300 kV and equipped with a Gatan US1000 CCD camera (2k × 2k). Images were captured every 1.0° over a ±60° range. The IMOD software package (http://bio3d.colourado.edu/imod) was used for the calculation of electron tomograms (Kremer et al., 1996). The Amira software package with an extension to the filament editor was used for the segmentation, automatic tracing, stitching and 3D visualization of microtubules, and for data analysis (Stalling et al., 2005; Weber et al., 2012; Weber et al., 2014).

### Spinning disk confocal fluorescence imaging

Live imaging was performed using a spinning disk confocal microscope (Nikon Ti2000, Yokugawa CSU-X1), equipped with 488-nm and 561-nm diode lasers, an EMCCD camera (Hamamatsu), and a 60X water-immersion objective (CFI Plan Apo VC 60X WI, NA 1.2, Nikon). Acquisition parameters were controlled by a home-developed LabVIEW program (LabVIEW, National Instruments). For *C. elegans* mitotic and female meiotic spindles, images were acquired every 2 second or 4 second with a single z-plane. For Human mitotic spindles, images were acquired every 2 second or 4 second with 3 z-sections every 1 μm, and the middle planes were presented.

### Laser ablation

The laser ablation system was constructed on the above-mentioned spinning disk confocal microscope. Femtosecond pulsed near-infrared lasers with either 80-MHz or 16-kHz repetition rate were adapted to perform laser ablation. 80-MHz femtosecond pulses with 0.3-nJ pulse energy and 800-nm center wavelength came directly from a Ti:sapphire pulsed laser (Mai-Tai, Spectra-Physics, Mountain View, CA). A 16-kHz femtosecond pulse train with ~6-nJ pulse energy was produced by selecting pulses from the above Ti:sapphire pulsed laser using a pulse picker (Eclipse Pulse Picker, KMLabs). The ablation laser was focused through the same objective for imaging, and laser ablation was performed by moving the sample on a piezo-stage (P-545 PInano XYZ, Physik Instrumente) in three dimensions controlled by a home-developed LabVIEW program (LabVIEW, National Instruments). Scanning line-cuts with z-steps were created by moving the stage perpendicular to the spindle long axis back and forth on the focal plane while lowering the stage in the z direction. The parameter for ablation in length by depth was 8 × 6 μm for *C. elegans* mitotic spindles; 12 × 6 μm for human mitotic spindles; 6 × 2 μm for C. elegans meiotic spindles. The moving speed of the stage was 50 μm/sec.

### Photobleaching experiments

Photobleaching experiments used the same set-up and software control as the laser ablation, except that a 80MHz Ti:sapphire pulsed laser was used (800nm wavelength, ~70 fs pulse-width, ~0.1-nJ pulse energy, Mai-Tai, Spectra-Physics, Mountain View, CA). The parameter for photobleaching in length by depth was 7 × 6 μm in *C. elegans* mitotic spindles, and 6 × 2 μm in C. elegans meiotic spindles. The moving speed of the stage was 50 μm/sec.

### Photoconversion experiments

Photoconversion experiments used the same set-up and software control as the laser ablation, except that a 405nm continuous-wave diode laser was used (Thorlabs, Inc.). The parameter for photoconversion in length was 12 μm in human mitotic spindles. The moving speed of the stage was 50 μm/sec.

### Quantitative analysis in *C. elegans* mitotic spindles

Quantitative analysis of the chromosome distance with a sub-pixel resolution was achieved using a home-written Matlab (The MathWorks, Natick MA) program. The center positions of two centrosomes in each frame of the time-lapsed images were either manually selected in spindles expressing GFP-labeled tubulin or automatically located in spindles expressing mCherry-labeled γ-tubulin based on a public Matlab program for particle tracking (https://site.physics.georgetown.edu/matlab/index.html). The center positions of the two centrosomes in each frame determines the spindle long axis as well as the spindle length, and were used to generate line-scans of overlaid images of mCherry-labeled histones (corresponding to chromosomes). Averaged mCherry fluorescence intensities from histones along these line-scans were extracted, and the fluorescence profile around histone-enriched regions were Gaussian-like shapes. A double-peak Gaussian function was used for fitting the line-scans to locate the centers of two groups of histones, presumably reflecting the central positions of sister chromosomes. The distance of these two fitted center positions was defined as the chromosome-to-chromosome distance. A straight line was fitted to the change of chromosome distance versus time to extract the velocity of chromosome separation before and after laser ablation. In controls, the velocity was computed before and after the chromosome distance was 2.4 μm, the chromosome distance around which most laser-ablation experiments were performed.

Movement and recovery of photobleaching makers were also tracked using a program written in Matlab. Line-scans of GFP-labeled tubulin between chromosomes along the spindle axis were extracted over the course of anaphase. Each line-scan at each time point can be divided into two halves by the middle plane of the spindle. The half of the line profile with the bleaching mark was normalized to the other half of the profile by a reflection of symmetry around the middle plane of the spindle. This profile normalization was used to remove spatial variations in the background fluorescence, a valid procedure assuming mirror symmetry of the spindle around its middle plane. A Gaussian function was used to fit the normalized profile to locate the center of the bleached mark, and thus the position of the bleached mark versus time was extracted. A straight line was fitted to the position of the bleached mark versus time to retrieve the velocity of the bleached mark.

### Quantitative analysis in *C. elegans* meiotic spindles

Chromosome distance in *C. elegans* meiotic spindles was computed with a combination of Fiji (Schindelin et al., 2012) and a Matlab program. Time-lapse images of spindle expressing mCherry::histone (corresponding to chromosomes) were realigned in a routine for matching, rotation and translation using Rigid Body of Fiji’s StackReg plug-in, so that the random displacement of the spindle due to the spontaneous motion of the worm was corrected. On the realigned time-lapse stack, a straight line passing through sister chromosomes was manually drawn in line with the spindle axis, and the kymograph of mCherry:histone intensities along this line was generated. Upon a Matlab program, each line-scan of mCherry:histone was fitted to a double-peak Gaussian function for computing the center positions of sister chromosomes, and thus the chromosome distance. A straight line was fitted to the data of chromosome distance versus time to extract the velocity of chromosome separation before and after laser ablation. In controls, separation velocity was computed before and after the chromosome distance was 3.3 μm, the chromosome distance around which most laser-ablation experiments performed.

Movement of photobleaching makers were also tracked using a program written in Matlab. Line-scans of GFP-labeled tubulin between chromosomes along the spindle axis were extracted over the course of anaphase. A Gaussian function was used to fit the line-scanning profile to locate the center of the bleached mark, and thus the position of the bleached mark versus time was extracted. A straight line was fitted to the position of the bleached mark versus time to retrieve the velocity of the bleached mark.

### Quantitative analysis in Human mitotic spindles

Distance information of interest was extracted from spindles expressing GFP::CENP-A (corresponding to kinetochores) and GFP::centrins (corresponding to spindle poles) in time-lapse z-stacks based on the following steps. First, using approaches from a particle tracking algorithm (Pelletier et al., 2009), a GUI Matlab program was developed to retrieve the three-dimensional coordinates of the center positions of each kinetochores and the two poles. Next, a support vector machine (SVM) algorithm was adapted to find the best plane to separate kinetochores of one half-spindle from those of the other half-spindle. This best plan is the one with the largest margin between two sides of kinetochores, and corresponds to the middle plane of the spindle. A unit vector perpendicular to this plane was thus in line with the direction of spindle long axis. All positions of kinetochores and spindle poles in space were projected onto this unit vector, and thus converted to one-dimensional information of projection lengths. Finally, chromosome-to-chromosome distance was computed as the length difference between the averaged projection lengths of kinetochores at one side of the half-spindle and those at the other side. Similarly, the chromosome-to-pole distance was computed as the length difference between averaged projection lengths of kinetochores and the projection length of the pole at the same side of the half-spindle. The above procedure was performed for each time point of a z-stack, so that the rotation and translation of spindles in space over the course of anaphase was corrected. A straight line was fitted to the data of chromosome-to-chromosome distance, and chromosome-to-pole distance versus time to compute the separation velocity of the chromosomes, and the velocity between chromosomes and poles before and after laser ablation, individually. In controls, the separation velocity of chromosomes was computed before and after the chromosome distance was at 6.1 μm.

Movement of photoconverted mEOS3.2-labeled microtubules were also tracked using a program written in Matlab. Line-scans of photoconverted tubulin between chromosomes along the spindle axis were extracted over the course of anaphase. A Gaussian function was used to fit the profile to locate the center of the photoconverted microtubules, and thus the position of the photoconverted microtubules versus time was extracted. A straight line was fitted to the position of the photoconverted microtubules versus time to retrieve the velocity of the photoconverted microtubules.

### Statistical analysis

Statistics are presented as mean ± SEM, and p-values were calculated by “ttest2” function in Matlab.

## Acknowledgements

We thank Marie Delattre for the *C. elegans* strain expressing GFP::tubulin and mCherry;tubulin, and the bacteria containing *gpr-1/2 (RNAi)* plasmid. We also thank Tobias Fürstenhaupt for technical assistance during acquisition of the tomographic data at the MPI-CBG (Dresden). We thank Marie Delattre and John Calarco for discussions. Work was supported by the Human Frontier Science Program (RGP 7 0034/2010) to D.N., Marie Delattre, and T.M.R. Work of D.N. and C.H.Y. was supported by the National Science Foundation grant DMR-0820484 and the National Institutes of Health Grant 1R01GM104976-01. T.M.R. received funding from the German Research Foundation (DFG grant MU 1423/8-1) and from the Saxonian State Ministry for Science and the Arts (SMWK). S.R. was funded by the Frauenhabilitationsförderung of the Faculty of Medicine Carl Gustav Carus of the TU Dresden. R.K. received funding from the European Union’s Horizon 2020 research and innovation programme under the Marie Skłodowska-Curie grant agreement No 675737 (grant to T.M.R.). The authors declare no competing financial interests.

## Author contributions

C.H.Y. and D.N. conceived and designed the research; C.H.Y. performed optical measurements and quantitative analysis; S.R., R.K., and T.M.R. performed electron tomography and 3D reconstructions. H.Y.W. and C.H.Y built the home-made laser ablation system. C.H.Y, H.Y.W. and R.F. prepared *C. elegans* samples; T.Y.Y. and C.H.Y. cloned and prepared human tissue culture cells. The manuscript was written by C.H.Y. and D.N., and commented by S.R. and T.M.R.

## Materials & Correspondence

Further information and requests for resources and reagents should be directed to and will be fulfilled by the corresponding author, Che-Hang Yu.

## Supplemental information

### Movie S1

Photoconversion (t=0 sec) of microtubules between chromosomes in a wildtype U2OS cell expressing GFP::CENP-A (green) and mEOS3.2::tubulin (green before photoconversion; magenta after photoconversion). Image size, 23 × 23 μm. This movie corresponds to Figure 2B.

### Movie S2

Laser ablation (t=0 sec) of microtubules between chromosomes in a wildtype U2OS cell expressing GFP::CENP-A (green) and mCherry::tubulin (magenta). Image size, 23 × 19 μm. This movie corresponds to Figure 3A.

### Movie S3

Chromosome segregation of a wildtype *C. elegans* embryo during anaphase. Chromosomes are colored in magenta; microtubules are colored in green. Image size, 48 × 36 μm. This movie corresponds to Figure 4A.

### Movie S4

Chromosome segregation of a *gpr-1/2 (RNAi) C. elegans* embryo during anaphase. Chromosomes are colored in magenta; microtubules are colored in green. Image size, 48 × 36 μm. This movie corresponds to Figure 4A.

### Movie S5

Laser ablation (t=0 sec) of microtubules (green) between chromosomes (magenta) in a *gpr-1/2 (RNAi) C. elegans* embryo. Image size, 30 × 13 μm. This movie corresponds to Figure 5A.

### Movie S6

Laser ablation (t=18 sec) of microtubules (green) between chromosomes (magenta) and poles (magenta) on both sides of the spindle in a wildtype *C. elegans* embryo. Image size, 45 × 45 μm. This movie corresponds to Figure 5B.

### Movie S7

Laser ablation (t=0 sec) of microtubules (green) between chromosomes (magenta) in a wildtype *C. elegans* ooctye in uterus. Image size, 22 × 22 μm. This movie corresponds to Figure 6B.

